# Milk of cow and goat, immunized by recombinant protein vaccine ZF-UZ-VAC2001(Zifivax), contains neutralizing antibodies against SARS-CoV-2 and remains active after standard milk pasteurization

**DOI:** 10.1101/2022.02.14.480298

**Authors:** Victoria Garib, Stefani Katsamaki, Shahlo Turdikulova, Yuliya Levitskaya, Nodira Zahidova, Galina Bus, Kristina Karamova, Manona Rakhmedova, Nigora Magbulova, Alexander Bruhov, Firuz Y. Garib, Ibrokhim Y. Abdurakhmonov

## Abstract

Here we report the first experimental validation of the possibility for obtaining immune milk with neutralizing antibodies against SARS-CoV-2 from vaccinated cow and goat using recombinant protein human vaccine, ZF-UZ-VAC2001. In the period of two weeks after first vaccination, we detected the neutralizing antibodies against coronavirus in the blood serum of vaccinated animals. The neutralizing activity, in its peak on the 21st days after receiving the third dose (77th day from first dose), was effective in Neutralization Test using a live SARS-CoV-2 in Vero E6 cells, even after 120-fold serum titration. Colostrum of the first day after 3rd dose vaccinated cow after calving had a greater activity to neutralize the SARS-CoV-2 compared to colostrum of subsequent three days (4.080 µg/ml vs 2.106, 1.960 and 1.126 µg/ml), goat milk (1,486 µg/ml), and cow milk (0.222 µg/ml) in MAGLUMI® SARS-CoV-2 neutralizing antibody competitive chemiluminescence immunoassay. We observed a positive correlation of receptor-binding domain (RBD)-specific IgG antibodies between the serum of actively immunized cow and milk-feeding calf during the entire course of vaccination (r = 0.95, p = 0.027). We showed an optimal regime for immune milk pasteurization at 62.5°C for 30 min, which retained specific neutralizing activity to SARS-CoV-2, potentially useful for passive immunization against coronavirus infection threats.

## Dear Editor

In the global coronavirus disease pandemic, immunological studies proved that antibodies are the effective molecules for sanitizing the body from viruses. However, the formation of novel SARS-CoV-2 mutations is leading to a decreased effectiveness of vaccines. Moreover, the slow rate of massive vaccination process, due to poor public acceptance and/or insufficient vaccine supplies in some countries, are the one of the main factors for continuous reemergence of new virus variants of concern (VOC). The development and registration of new vaccines against constantly emerging mutations require additional time and funding. This underlies to explore new opportunities to establish a stable herd immunity, focusing on the development of highly effective neutralizing antibodies (nAbs) [1]. Apparently, nAbs against VOC can be quickly obtained by the vaccination of farm animals with the emergency use approved (EUA) human vaccines, covering the new mutations of importance. Here, we are to briefly communicate the first experimental validation report on obtaining immune milk with neutralizing antibodies (nAbs) against SARS-CoV-2 from cow and goat after vaccination with RBD-based recombinant protein subunit human vaccine, ZF-UZ-VAC2001 or (ZF2001 or Zifivax).

The idea, to study and validate if vaccinated farm animal milk contains nAbs or not, came from the evidence that antibodies to SARS-CoV-2 were found in the milk of women who have had COVID-19 or been vaccinated [2]. We also carefully read and acknowledge several publications on the possible benefit from the passive immunization using milk of vaccinated cow. Jawhara [3] first suggested that microfiltered raw immune milk or colostrum collected from SARS-CoV-2 vaccinated cows could provide short-term protection against SARS-CoV-2 infection in humans. Further, Arenas et al. [4] proposed to use of heterologous passive immunity using Bovine Coronavirus (BCoV) immune milk as an immunostimulant therapy to control SARS-CoV-2 infection because vaccination of farm animals is a well-known and has been described in the literature to protect animals from diseases, including BCoV [5]. Gallo et al. [6] reviewed the antiviral properties of native and chemically modified whey proteins and their potential applications in human health, focusing on their application in prevention and treatment of SARS-CoV-2 infection.

However, neither the detection of nAbs in the serum and milk of vaccinated household farm animals using human vaccines approved for emergency use, nor the effect of pasteurization of such immune milk on the SARS-CoV-2 virus-neutralizing activity, has not yet been experimentally validated. To address these limitations, we first performed a series of pilot experiments in 60 milk and serum samples of vaccinated household cows and goat. In that, healthy pregnant as well as dairy cow with feeding calf were vaccinated with the ZF-UZ-VAC2001 recombinant SARS-CoV-2 human vaccine, containing a dimeric form of the receptor-binding domain (RBD) as the antigen [7].

We used this particular vaccine because it was successfully passed a 3rd phase clinical trial in Uzbekistan and were approved for the massive vaccination of the population [8]. ZF-UZ-VAC2001 has shown its effectiveness against the diverged variants of the SARS CoV-2 virus [8,9], namely Alfa (92.93%), Gamma (100%), Delta (77.47%), and Kappa (90.0%). Although the neutralization activity of currently used human vaccines, has shown a decreased effect against emerging new VOC B.1.1.529 (Omicron) variant [9], the prolonged time between the second and third dose injections of ZF-UZ-VAC2001 (0,1, 2 vs 0, 1, 5 months) has revealed a better neutralization effect against Omicron variant [10].

In our study, we performed vaccination of farm animals (cows and goat) with revaccination carried after 28 and 56 days. Examination of the udder, palpation of the lymph nodes, mucous membranes and measurement of rectal temperature as well as the observation of the behavior of animals have not revealed any side effects during daily monitoring of animals after vaccination.

Since the first dose vaccine injection period, we have regularly collected and evaluated the virus neutralization activity of the immune milk and blood serum of the cow and goat, measuring the inhibition level of viral RBD binding to the ACE2 receptor. For this purpose, we used the MAGLUMI® SARS-CoV-2 Neutralizing Antibody competitive chemiluminescence immunoassay (CLIA). Milk and blood serum from the non-vaccinated cows were used as control samples. For blood serum samples, we performed the direct neutralization test using a live SARS-CoV-2 proliferating in Vero E6 cells [S1] at the Medical University of Vienna, Austria. We also determined antigen-specific antibody isotypes to the RBD of spike (S) and full-length S proteins in bovine serum and milk samples using enzyme-linked immunosorbent assay (ELISA) analysis (see Supplementary Materials and Methods).

We detected the neutralizing antibodies to the RBD domain and S-protein of SARS-CoV-2 in the blood serum of a vaccinated animal as early as two weeks after the first vaccination. Results showed that revaccination contributed to an increase in the effect of inhibition. We noted maximum neutralization activity of blood serum (100%) and milk (40%) on the 77th day from the date of the first vaccination. The correlation between neutralization rate of cow sera and milk during the entire course of vaccination was significant at r = 0.96, p = 0.022

Complete viral neutralization of the cytopathic effect on Vero E6 cells was detected even with 120-fold serum titration. The blood serum and milk of the vaccinated cow contained specific IgG to the RBD domain and S-protein of SARS-CoV-2.

We also found statistically significant correlation between the level of IgG specific to the RBD domain and the neutralization rate of milk of vaccinated cows (r = 1.0, p = 0.001, Supplementary Table 1).

When we vaccinated other group of “Simmental” cows during the dry period of the third trimester of pregnancy, after calving we determined that that the colostrum of the first day had a greater ability to neutralize the SARS-CoV-2 virus (4.080 µg/ml) compared to colostrum of subsequent next days (2.106, 1.960 and 1.126 µg/ml) and milk [0.222 µg/ml that was within the range of 50% (or 0.30 µg/ml) neutralization activity].

In another pilot experiment in a domestic lactating goat (*Capra hircus*), it was sufficient to use only 2 dose injection of the vaccine to achieve the effect of immunization and obtain immune milk samples. The correlation between neutralization rate of goat sera and goat milk in during the entire course of vaccination was high at r = 0.99, p = 0,003 (Supplementary Table 1). The maximum of virus-neutralizing activity of the immune goat milk was 1.486 µg/ml on the 14 days of the third dose vaccine injection.

We further investigated the possibility of transferring of SARS-CoV-2 specific IgG antibodies via milk from cow to its milking calf. Toward this goal, we performed the analysis of sera from calf during active vaccination of cow. Results showed that there is statistically significant correlation (r = 0.95, p = 0.027, Supplementary Table 1) between the level of IgG specific to RBD of vaccinated cow and calf fed by milk of vaccinated mother cow. This showed the possibility of passive immunization using milk of vaccinated mother cows, containing specific IgG against SARS-CoV-2.

In perspectives, as highlighted above [2-6], there is a huge opportunity to use the passive immunization property of immunized cow milk in humans. For this, there is a need for determining the optimal pasteurization conditions of immune milk to keep its biologically active neutralization against SARS-CoV-2. Toward this goal, we have studied several pasteurization regimes of immunized milk while analyzing its neutralization activity of SARS-CoV-2. We tested 20 milk samples obtained from vaccinated cow and goat, pasteurizing them at different temperatures and regimes of pasteurization according to State regulations (62.5-63°C for 30 min, 72°C for 5 min, 85°C for 5 sec).

As expected, temperature treatment for pasteurization of milk product negatively correlated with neutralization activity (r =-1.0, p = 0.02). Compared to before-pasteurization raw milk (0.5±0.2 µg/ml), pasteurization treatments at 72 ºC (for 5 min) and 85 ºC (for 5 sec) temperatures showed sharp activity decreases (0.1±0.04 µg/ml and 0.05±0.01 µg/ml respectively). However, the pasteurization at 62.5-63 ºC during 30 minutes retained sufficiently active neutralization property in nAbs chemiluminescence immunoassay (0.3±0.1 µg/ml), small decrease observed between raw and pasteurized immune milk at 62.5-63 ºC during 30 minutes was statistically non-significant (p = 0.24; (Fig.1; Supplementary Table 2).

**Fig. 1.**
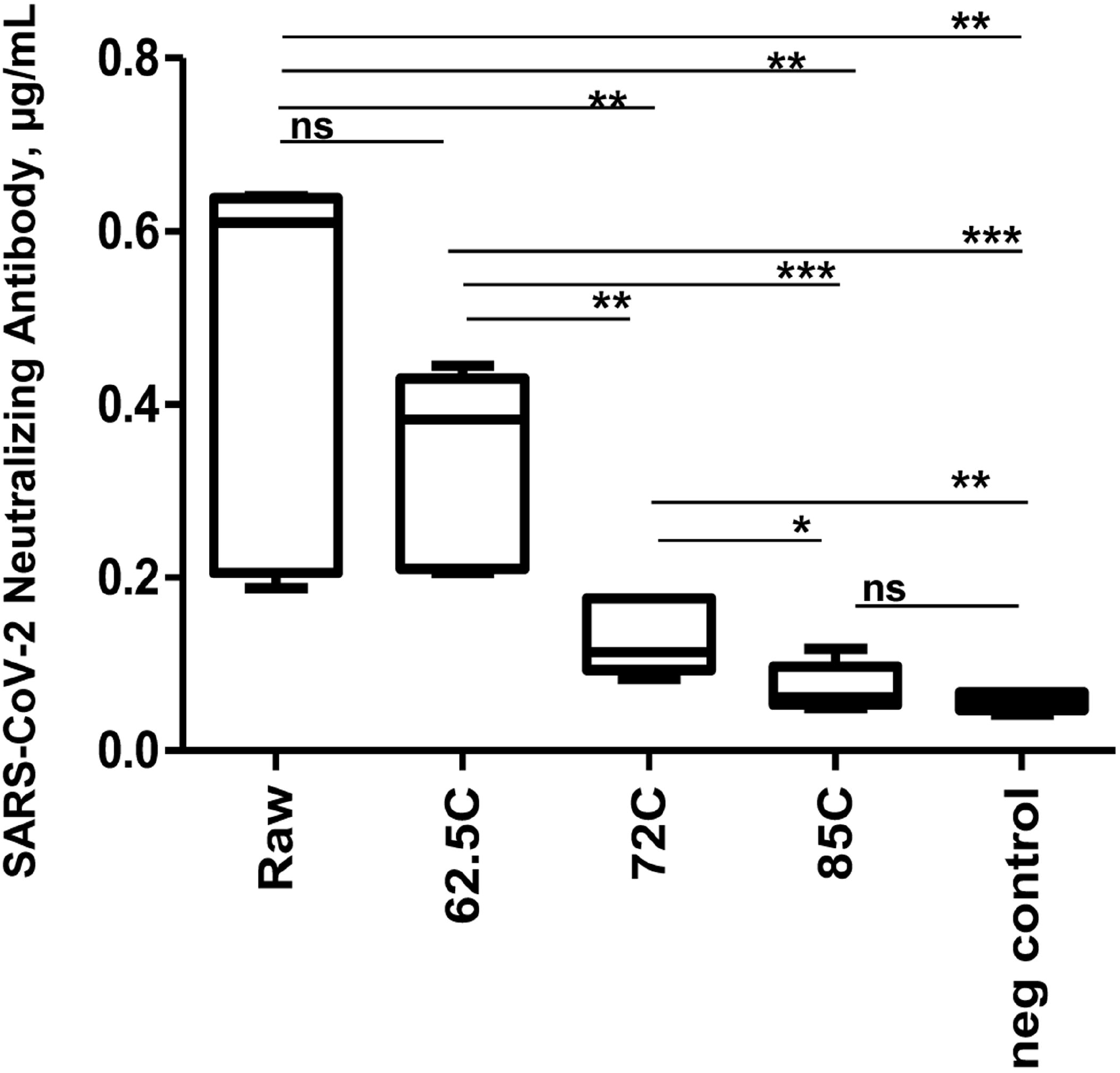
The box plots of SARS-CoV-2 Neutralizing Antibody levels (µg/mL). Average indices (y-axis) for before pasteurization (raw) and pasteurized immune milk samples at the different regimes of pasteurization (x-axis) are shown. *p ≤ 0.05, **p ≤ 0.01, ***p ≤ 0.01; ns: not statistically significant difference. Negative control represents milk sample from non-vaccinated animal. The box plots were generated, using GraphPad Prism 8 (GraphPad Software Inc.).

Thus, these first experimental validation results indicated that the ZF-UZ-VAC2001 human vaccine can be safely used for the vaccination of household farm animals to obtain immuno-biologically active milk and serum, containing specific nAbs against SARS-CoV-2. There is no need for the substantial corrections in dose calculations for vaccination of the animals, which can be performed in accordance to the manufacturer’s instructions and protocols for human vaccination. Interestingly, we experimentally observed that milk and serum of vaccinated cow not only have active neutralization property against SARS-CoV-2, but also biologically active specific IgG antibodies to RBD and S protein, transferred from mother cow milk to its milking calf, passively forming offspring’s immune protection.

We found that the most optimal condition for immunized milk pasteurization can be achieved 62.5-63 ºC during 30 minutes, which retains efficient virus neutralizing antibody activity against SARS-CoV, providing future opportunity for the passive immunization of human by milk consumption. The limitation of this study is a small number of farm animals included for vaccination experiments and future large-scale experiments are required, which is in progress in our laboratories. However, early results collectively showed that pasteurized immune milk obtained from vaccinated household animals using human vaccine against SARS-CoV-2, administered herein, potentially could be useful for passive immunization against coronavirus infection threats. The use of immune colostrum, milk and dairy products with neutralizing antibodies from vaccinated cows and goats seems to be a promising approach for the preparation of safe and natural prophylactic agents against human and animal infections.

## Abbreviations

ACE2: angiotensin-converting enzyme 2
COVID-19: coronavirus disease
ELISA: enzyme-linked immunosorbent assay
HRP: horseradish peroxidase
IgG, IgA, IgM: immunoglobulin G, A, M
nAbs: neutralizing antibodies
NT: Neutralization test
OD: optical density
RBD: receptor-binding domain
S: spike protein
S1: spike protein receptor-binding subunit
SARS: severe acute respiratory syndrome
VOC: variants of concern
CLIA: chemiluminescence immunoassay
EUA: emergency use approval
ZF-UZ-VAC2001: Zhifei (ZF) – Uzbekistan (UZ) –Vaccine (VAC) project started in January of 2020 (20/01)

## Supplementary Information

The online version contains supplementary material available at https://doi.org/xxx

**Supplementary Table 1**. Statistical analysis of correlations between parameters.

**Supplementary Table 2**. P-values of significant difference between nAbs levels of immune milk at different pasteurization (n=20).

**Supplementary Materials and Methods**.

## Acknowledgments

The authors thank Assoc. Prof. Priv.-Doz. Dr. Karin Stiasny and Ing. Jutta Hutecek from Centre of Virology, Medical University of Vienna, Austria for performing the SARS-CoV-2 Neutralization Test, group of Prof. Rudolf Valenta from Department of Pathophysiology and Allergy Research, Medical University of Vienna, Austria for fruitful collaboration. We thank also specialists and workers of the “Panaev’s Animal Farm” Karakalpakistan of Republic of Uzbekistan for their help and support of the study. We acknowledge the collaborative efforts of the Anhui Zhifei Longcom Biopharmaceutical, Hefei, China as well as CAS Key Laboratory of Pathogenic Microbiology and Immunology, Institute of Microbiology, Chinese Academy of Sciences, Beijing, China on the vaccine development project.

## Authors’ contributions

**VG** and **IYA**: project administration, conceptualization, methodology, investigation, data curation, and writing—original draft, review and editing; **FYG**: consultation, methodology, writing—review and editing; **YL** and **ST**: methodology, investigation, writing—review and editing; **SK GB, KK, MR, NM**, and **AB**: investigation, data collection and curation. All authors contributed to the article and approved the submitted version.

## Availability of data and materials

The datasets used and/or analyzed during the current study are available from the corresponding author on a reasonable request.

## Declarations

### Ethics approval and consent to participate

The studies reviewed and approved by the Ethics Committee Republic of Uzbekistan for SARS-CoV-2 research (authorization number 6/6 1449 from 13.10.2020)

### Consent for publication

All authors consent to the publication of the manuscript in Nutrition Journal.

