## Supplementary Materials and Methods for "Milk of cow and goat, immunized by recombinant protein vaccine ZF-UZ-VAC2001(Zifivax), contains neutralizing antibodies against SARS-CoV-2 and remains active after standard milk pasteurization"

**Animal vaccination**

Farm animals were kept separately in “Panaev’s Animal Farms” conditions, Karakalpakistan, Republic of Uzbekistan. Animals were vaccinated with recombinant ZF-UZ-VAC-2001 human vaccine against SARS-CoV-2 virus (also named ZF2001 or Zifivax). ZF-UZ-VAC-2001 is a protein subunit human vaccine using a dimeric form of the receptor-binding domain (RBD) as the antigen, collaboratively developed by Anhui Zhifei Longcom [7]. Revaccination was carried out intramuscularly after 28 and 56 days, according to the manufacturer's instructions by analogy with those recommended for use in humans, i.e., 1 dose per 70 kg of animal weight. The sampling of milk and blood was carried out in the morning. The studies reviewed and approved by the Ethics Committee of the Republic of Uzbekistan for SARS-CoV-2 research (authorization number 6/6 1449 from 10/13/2020).

**Preparation of milk and colostrum samples**

We used the protocol of Fox et al.[S2] for human milk to remove lipids from milk samples. Briefly, milk samples were centrifuged at 800g for 15 min at room temperature, fat was removed, and supernatant was transferred to a new tube. Centrifugation was repeated 2x to ensure removal of all cells and fat. Skimmed acellular milk was aliquoted and frozen at -80°C until testing. In the case of freezing, the samples were re-cleaned by centrifugation as above after thawing. To study the influence of pasteurization, three different milk samples were preliminarily heated using the various regimes (62,5-63°C for 30 min; 72°C for 5 min; and 85°C for 5 sec) according to State regulations for pasteurization (N 0281-09).

**Virus neutralization assays**

*MAGLUMI® SARS-CoV-2 Neutralizing Antibody assay (CLIA)* was performed using fully-auto chemiluminescence immunoassay analyzer MAGLUMI series, according to the manufacturer's instructions for human sera (Snibe Diagnostic, Pingshan, China) at the Scientific and Diagnostical Centre of Laboratory Technology "Defactum Laboratories", Tashkent, Uzbekistan. Briefly, milk sample was thoroughly mixed with magnetic microbeads coated with ACE2 antigen and recombinant SARS-CoV-2 S-RBD antigen labelled with ABEI. Mixed content was incubated in a buffered condition. SARS-CoV-2 Neutralizing Antibody, present in milk samples, competes with ACE2 antigen immobilized on magnetic microbeads for binding recombinant SARS-CoV-2 S-RBD antigen labelled with ABEI. After precipitation in a magnetic field, the supernatant was decanted and pelleted magnetic beads were washed. Subsequently, the Starter 1+2 reagent was added to initiate a chemiluminescent reaction. The light signal was measured by a photomultiplier as relative light units (RLUs), which is inversely proportional to the concentration of SARS-CoV-2 neutralizing antibody presented in the milk sample. The analyzer automatically calculated the concentration in each sample by means of a calibration curve, which was generated by a 2-point calibration master curve procedure. The results are expressed in μg/mL, where 0.300 μg/mL is considered a value of the sample that provide a 50 % inhibition of viral growth.

*SARS-CoV-2 Neutralization test (NT)* was carried out in a laboratory with a BSL3 security level of the Virology Centre of the Medical University of Vienna, Austria according to the protocol Koblischke et al. [S1]. Briefly, two-fold serial dilutions of heat-inactivated samples were incubated with 50–100 TCID50 SARS-CoV-2 for1 h at 37◦C before the mixture was added to Vero E6 cell monolayers (starting dilution of samples 1:10). Incubation was continued for 2–3 days. NT titers were expressed as the reciprocal of the sample dilution required for 100% protection against virus-induced cytopathic effects. NT titer ≥10 was considered positive.

**Determination of bovine anti-RBD- and anti-S-protein specific IgG, IgA, and IgM**

The determination of antigen-specific antibody isotypes to the receptor-binding domain (RBD) of spike (S) protein and full-length S protein in bovine serum and milk samples by enzyme-linked immunosorbent assay (ELISA). Aliquots of 2 µg/ml of receptor-binding domain (RBD) of spike (S) protein or full-length S protein (Genscript, Leiden, Netherlands) were coated overnight onto NUNC Maxisorp 96 well plates (Thermofisher Scientific, Massachusetts, USA). After three times washing with washing PBST buffer and blocking with 2% BSA (bovine serum albumin) solution, plates were incubated at 4° overnight. Serum or milk dilutions (1:500 / 1:50 / 1:10 in 0,5% BSA) in amount 100 µl/well were added after removing of blocking solution and stored overnight at 4°C.

The plates were washed three times with 200 µl PBST and 50 µl of peroxidase-labelled sheep anti-bovine IgG, IgM, or IgA Ab (Bio Rad, Hercules, California, USA) diluted in dilution buffer 1:25000 was added per well and incubated at RT for 120 min with flowing five times washing steps. The colour reaction was started by adding 50 µl/well of substrate solution (200 mg 2,2’-azino-bis (3-ethylbenzothiazoline-6-sulphonic acid (ABTS; Sigma-Aldrich, Darmstadt, Germany); in 200 mL citric buffer (61.5 mM citric acid, 77.3 mM Na_2_HPO_4_ × 2 H_2_O, pH 4) and 20 µL hydrogen peroxide). The optical density (OD) values corresponding to the levels of antigen-specific antibodies were determined at 405 and 492 nm in an TECAN Infinite F5 ELISA reader. All determinations were performed in duplicates and results are shown as mean values with a variation of <5%.

**Data analyses and statistical analysis**

Data analysis and statistical significance of correlations were performed using Pearson Correlation - Free Statistics Software (Calculator) [S3]. We computed Pearson Correlation including Pearson Product Moment Correlation, Covariance, Determination, and the Correlation T-Test determining statistically significance from both one- and two-sided p-values. The Jarque-Bera and Anderson-Darling Normality Tests were applied to both variables. Significant differences between groups were calculated using one-way analysis of variance (ANOVA) followed by Bonferroni’s posttest. Statistical significance was determined at p ≤ 0.05 - 0.001 levels (Supplementary Tables 1 and 2). The box plots were generated, using GraphPad Prism 8 (GraphPad Software Inc.).

**Supplementary Table 1.** Statistical analysis of correlations between parameters.

| **Parameters** | **Covariance** | **Correlation** | **Determination** | **T-Test** | **p-value (2 sided)** | **p-value (1 sided)** |
| --- | --- | --- | --- | --- | --- | --- |
| Neutralization rate of cow sera vs milk in during vaccination | 744,25 | 0,96 | 0,91 | 4,59 | 0,044 | 0,022 |
| IgG-RBD vs neutralization rate of milk | 15,86 | 1,00 | 1,00 | 24,35 | 0,002 | 0,001 |
| IgG-RBD of sera after active vs passive immunization | 0,31 | 0,95 | 0,90 | 4,16 | 0,053 | 0,027 |
| Temperature of milk pasteurization vs neutralization rate | -93,83 | -1,00 | 1,00 | -14,13 | 0,045 | 0,022 |
| Neutralization rate of goat sera vs milk in during vaccination | 9,54 | 0,99 | 0,99 | 12,53 | 0,006 | 0,003 |
| Neutralization rate of cow milk vs goat milk in during vaccination | 0,05 | 0,98 | 0,96 | 6,54 | 0,023 | 0,011 |

**Supplementary Table 2.** P-values of significant difference between nAbs levels of immune milk at different pasteurization (n=20).

| sample | raw | 62.5C | 72C | 85C | neg control |
| --- | --- | --- | --- | --- | --- |
| raw | – | 0.2405 | 0.0069 | 0.0026 | 0.0020 |
| 62.5C | 0.2405 | – | 0.0030 | 0.0004 | 0.0003 |
| 72C | 0.0069 | 0.0030 | – | 0.0139 | 0.0035 |
| 85C | 0.0026 | 0.0004 | 0.0139 | – | 0.2903 |
| neg control | 0.0020 | 0.0003 | 0.0035 | 0.2903 | – |
